# Public goods exploitation is reduced in species-rich microbial communities

**DOI:** 10.1101/2022.11.25.517952

**Authors:** Siobhán O’Brien, Chris Culbert, Timothy G. Barraclough

## Abstract

Intraspecific public goods are commonly shared within microbial populations, where the benefits of public goods are largely limited to closely related conspecifics. One example is the production of iron-scavenging siderophores that deliver iron to cells via specific cell envelope receptor and transport systems. Intraspecific social exploitation of siderophore producers is common, since non-producers avoid the costs of production but retain the cell envelope machinery for siderophore uptake. However, little is known about how interactions between species (i.e. interspecific interactions) can shape intraspecific public goods exploitation. Here, we predicted that strong competition for iron between species in diverse communities will increase costs of siderophore cooperation, and hence select for increased intraspecific exploitation. We examined how increasing microbial community species diversity shapes intraspecific social dynamics by monitoring the growth of siderophore producers and non-producers of the plant-growth promoting bacterium *Pseudomonas fluorescens*, embedded within tree-hole microbial communities ranging from 2-15 species. We find, contrary to our prediction, that siderophore exploitation is reduced in high diversity communities, driven by increased likelihood of encountering key species that reduce the growth of siderophore non-producing (but not producing) strains of *P. fluorescens*. Our results suggest that maintaining a healthy soil microbiota could contribute to the maintenance of siderophore production in natural communities.

## 1. Introduction

Microbes exhibit a wide range of cooperative behaviours that can shape, and be shaped by, the communities within which they reside [1, 2]. Siderophore production by bacteria and fungi is one well-studied example of a cooperative behaviour [3–5]. Under iron-limitation, extracellular siderophores are produced that deliver iron to the cell via specific receptor and transport systems [6]. However, since siderophore production is metabolically costly, it can be exploited by non-producing cheats who avoid the cost of production while benefiting from siderophores of nearby cooperators [3]. Such non-producers invade populations of cooperators under iron-limited conditions and can evolve *de novo* within days [7, 8].

Selection for non-producing phenotypes depends on the relative costs and benefits of siderophore production. The cost of siderophore production increases as iron becomes limited and genes associated with siderophore production are upregulated [9]. When costs of cooperating are higher, there is increased selection for cheating, as non-producers experience a large relative fitness advantage by avoiding or reducing the cost of cooperating. Paradoxically, this means that cheating is strongly favoured when siderophores carry large benefits [9, 10]. Cooperators, on the other hand, will benefit from siderophore production when these benefits are more likely to accrue to the cooperators themselves or identical clonemates who also produce siderophores, for example via spatial heterogeneity or environmental viscosity [11, 12].

While the role of the abiotic environment is well-established in shaping the costs and benefits of siderophore production, less is known about whether the biotic environment (i.e., interactions between species) also alters the dynamics of within-species siderophore cooperation. Given that species can compete with one another for iron in natural ecosystems [13, 14], provide a source of iron upon cell lysis [15], or even pirate ‘foreign’ siderophores produced by other species [16], it is likely that species interactions can alter the costs and benefits of siderophore production in multiple, contrasting ways. For example, the presence of a second species (*Staphylococcus aureus*) can either promote or restrict the spread of *Pseudomonas aeruginosa* siderophore cheats by acting as a competitor or source of iron, respectively - depending on the degree of competition between the two species [15, 17]. However, our understanding of how species interactions alter siderophore production dynamics in natural communities remains largely limited to the aforementioned highly simplified two-species communities. In natural complex communities, a high degree of competition for resources such as iron [13, 14] or even the building blocks of siderophores themselves [18], could increase selection for non-producing cheats by increasing the cost of cooperation. This is important, as siderophore-producing bacteria are often relied upon for functions such as plant growth promotion and biocontrol, where they must interact with diverse soil, root or leaf microbiomes [19].

Here, we test whether increasing microbial community species diversity increases selection for cheats, by monitoring the growth of siderophore producers and non-producers of the plant-growth promoting bacterium *Pseudomonas fluorescens*, embedded within communities ranging from 2-15 species. We predicted that siderophore exploitation should be greatest in high diversity communities, where between-species competition is more intense. Community taxa originated from semi-permanent rain-filled wells formed by the roots of beech trees (*Fagus sylvatica*) [20], and competition has previously been reported to dominate between wild isolates of these taxa [21]. We find, contrary to our prediction, that siderophore exploitation is reduced in high diversity communities, suggesting that maintaining a healthy soil microbiota could contribute to the maintenance of siderophore production in natural communities.

## 2. Methods

### a) Focal species–*Pseudomonas fluorescens*

We used the gentamicin-resistant *lac-Z* marked *Pseudomonas fluorescens* strain SBW25-*lacZ* as our siderophore producing strain [22] and the gentamicin-resistant strain *SBW25ΔpvdL* which lacked genes encoding the primary siderophore pyoverdine, as our non-producing strain [23]. Gentamicin resistance distinguished our focal strains from the rest of the community (see below). *LacZ* conferred producers (SBW25-*lacZ*) a blue pigment, so that they could be easily distinguished from non-producers (SBW25Δ*pvdL*) on Lysogeny Broth (LB) agar supplemented with 90 μg/mL 5-Bromo-4-chloro-3-indolyl-β-Dgalactopyranoside (X-gal). Previous work has shown that fitness levels of SBW25-*lacZ* are comparable to the wild-type ancestor, suggesting the cost of the *lacZ* marker is negligible or absent [22].

### b) Background microbial community

We screened 230 isolates from an archived tree-hole library [24] for i) gentamicin susceptibility (to permit differentiation from focal *P. fluorescens* strains) and ii) growth in iron-limited lysogeny broth (LB) (to ensure survival under the experimental conditions). Gentamicin susceptibility of each isolate was tested using 10μg gentamicin antimicrobial susceptibility discs (Thermo Scientific^™^ Oxoid^™^). Growth in iron-limited LB was verified by measuring OD600 (600nm) of each isolate after 48h at 22^O^C in iron-limited LB (LB supplemented with 100 *μ*g/mL human apo-transferrin, an iron chelator, and 20 mM sodium bicarbonate [25]). This screening process produced a library of fifteen isolates from which we built our background communities. The identities of the chosen 15 isolates (Table 1) were confirmed through 16S rRNA sequencing (supplementary methods).

**Table 1:** Bacterial genera used in this study

| Strain | Genus | Experimental number |
| --- | --- | --- |
| BB19.36 | *Yersinia* | **1** |
| SP01.03 | *Acinetobacter* | **2** |
| SP01.04 | *Bacillus* | **3** |
| SP03.13 | *Janthinobacter* | **4** |
| SP06.03 | *Erwinia* | **5** |
| SP03.19 | *Epilithonimonas* | **6** |
| WYM27.02 | *Pantoea* | **7** |
| WYM29.03 | *Rhodococcus* | **8** |
| WYC41.02 | *Pseudomonas* | **9** |
| BB66.01 | *Aeromonas* | **10** |
| SP03.21 | *Serratia* | **11** |
| SP04.06 | *Pedobacter* | **12** |
| AE95.04 | *Stenotrophomonas* | **13** |
| BB19.34 | *Chryseobacter* | **14** |
| SP06.07 | *Buttiauxella* | **15** |

### c) Experimental design

We tested whether the growth of our focal *P. fluorescens* producer and non-producer was affected by the richness of the background community. We chose four levels of community richness: 2, 4,8, and 15. Each richness level includes the addition of *P. fluorescens*, for example, a community richness of 4 represents *P. fluorescens* plus 3 other species. We adopted a random partitions design (see [26]). Each richness level was represented by 5 random combinations of the 15 bacterial isolates, except when richness was equal to two; when equal to two, all 15 background isolates were grown with SBW25. Therefore, the experiment included 15 different two-species communities, 5 different four-species communities, 5 different eight-species communities and 5 different fifteen-species communities (Supplementary Table S1). Each community was replicated five times.

To test whether our focal *P. fluorescens* non-producer could exploit siderophores of the *P. fluorescens* producer under different levels of community diversity, we grew our focal *P. fluorescens* strains as single genotypes (producer and non-producer grown separately) and as mixed genotypes (producer and non-producer grown together in a 1:1 mixture). Including a mixed versus single genotype treatment allowed us to confirm that between-species siderophore exploitation was occurring in communities. Initial densities of our focal *P. fluorescens* species were always the same between single and mixed genotype conditions (*i.e*. as mixed genotypes, the starting densities of each strain was halved compared to single genotypes). Individual communities were assembled so all taxa within a community were inoculated at equal starting densities. Using a liquid handling robot (Hamilton Microlab STARlet), 20 *μ*l of bacterial cultures in total were added to 180 *μ*l of iron-limited LB media (prepared as described above) in 96 well microplates (Thermo Fisher Scientific) to a final volume of 200 *μ*l. The final cell density was always 400 cells/well. Sterile water was added to iron-limited LB in six wells of each plate as negative controls. Plates were incubated at room temperature for 7 days as batch cultures. Final cell densities of focal *P. fluorescens* were measured by plating on gentamicin-supplemented LB agar and counting colony forming units (CFU / ML).

### d) Statistical analysis

We estimated the total final population density (CFU / ML) for each *P. fluorescens* strain (producer or non-producer) in each community under both single and mixed genotype conditions. For single genotype conditions, we tested whether species richness (fitted as a continuous variable) and/or *P. fluorescens* genotype (producer or non-producer) affected final densities of *P. fluorescens*, using generalised mixed effects models (glmer), controlling for the identity of the background community as a random factor. The response variable (final densities) was log transformed to comply with model assumptions of normality. For mixed genotype conditions, we assigned proportion of non-producers (non-producer density / total *P. fluorescens* density) in each community as our response variable, with species richness as a continuous explanatory variable in a glmer with a binomial error structure and community as a random factor. We quantified the effect of each species on final densities of producers and non-producers as single genotypes using the linear model method described in [26], which includes presence or absence of each species as an explanatory variable in the model. The species coefficients provided by this method give a measure of the effect of each species on focal strain final densities relative to an average species. We analysed data using R Studio version 1.4.

## 3. Results

### a) Evidence for exploitation within the focal species

By comparing the final proportion of *P. fluorescens* non-producers in single versus mixed genotype communities, we find that mean proportion of non-producers increased when in co-culture with the producer (effect of genotype, *X*^2^_1,3_= 35.87, p<0.0001). This was irrespective of species richness level (genotype x richness, *X*^2^_1,4_= 0.29, p=0.58, Fig S1). In other words, non-producers reached higher frequencies in communities when their conspecific producer was also present – suggesting that within-species siderophore exploitation occurs in this system.

### b) *P. fluorescens* siderophore producer and non-producer growth increases and decreases respectively as communities become more diverse

#### (i) Single genotype condition

Under single genotype conditions (where the producer and non-producer are grown separately) species richness had opposing effects on our focal producer and non-producer final densities. Specifically, final densities of producers increased with species richness, whereas final cell densities of non-producers decreased as species richness increased (genotype x richness interaction, *X*^21,5^= 14.493, p=0.0001, Fig 1, Fig S2) This was mainly driven by significantly greater (173%) non-producer versus producer final densities in two-species communities (p<0.001, tukey post-hoc test, Fig 1, Fig S2). The advantage of non-producers over producers in two-species communities was reduced in communities comprising 4, 8 and 15 species, where there was no difference between producer and non-producer final cell densities (p > 0.5 in 4, 8 and 15 species communities, tukey post-hoc tests, Fig 1, Fig S2).

**Fig 1:**
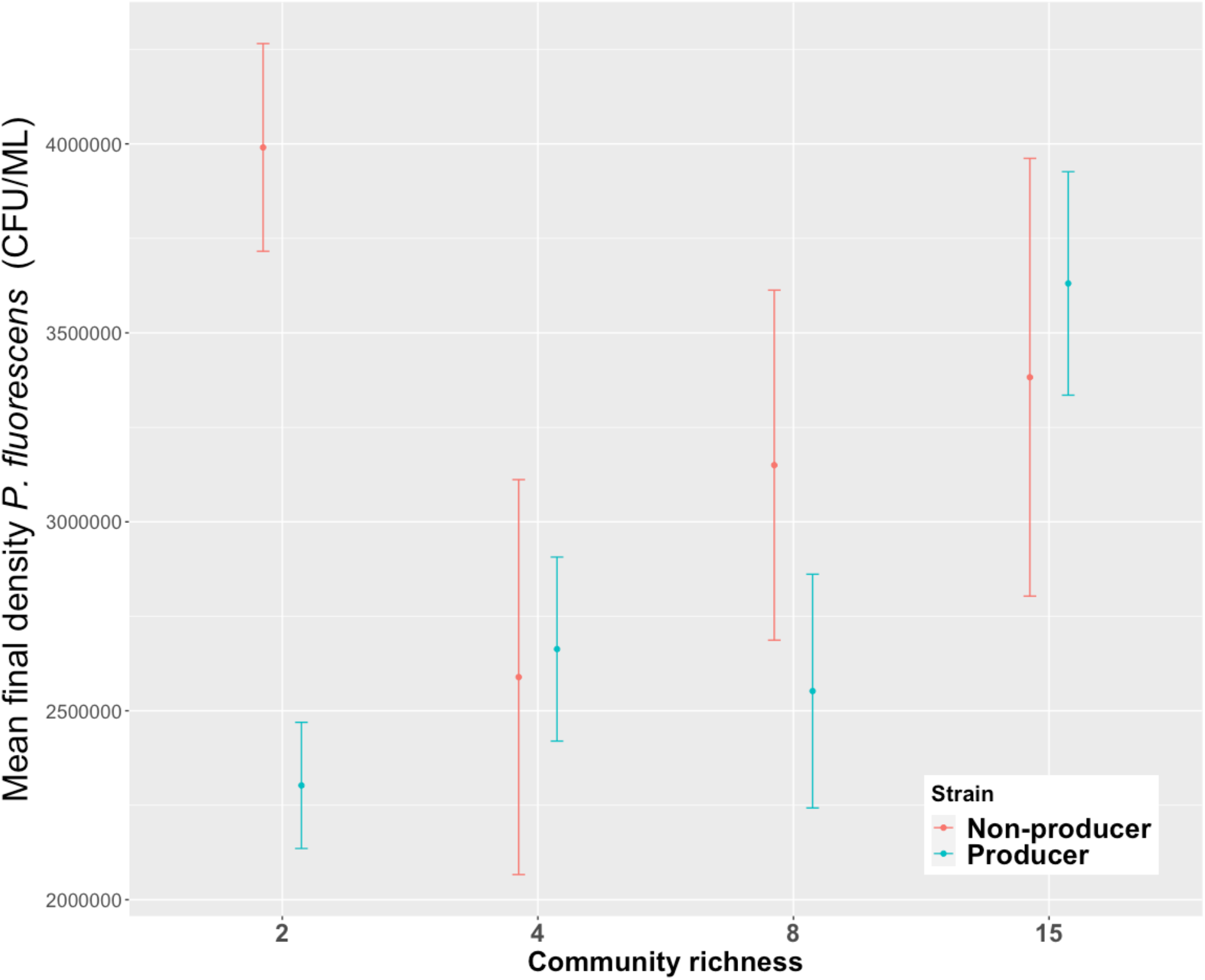
Mean final densities (CFU/ML) of single genotype *P. fluorescens* producer and non-producer populations in communities comprising 2, 4, 8 and 15 species. We find that species richness has opposing effects on our focal producer and non-producer final densities when inoculated as single genotypes. Producers grew better in high richness compared to low richness communities, while non-producers grew better in low richness compared to high richness communities (genotype x richness interaction, *X*^2^_1,5_= 14.493, p=0.0001). Sample size varies between richness levels. When r=2, n=75. When r=4,8 or 15, n=25. Points show mean value ±SE.

#### (ii) Mixed genotype condition

Given that siderophore producers reach higher densities in complex versus simple communities, we next tested whether increased densities of producers would paradoxically allow non-producers to reach higher frequencies in complex versus simple communities. Contrary to our prediction, we found that when both producer and non-producer genotypes were grown together (mixed genotype treatment), the proportion of producers in the community increased with species richness (*X*^2^_1,2_= 10.27, p=0.001, Fig. 2). This is in line with our single genotype result (above) indicating that producer and non-producer final densities reduced and increased, respectively, as communities became more diverse. Importantly, these findings further show that while increased community complexity can contribute to the maintenance of siderophore production, this was independent of intraspecific exploitation (which could not occur in the single genotype treatment where non-producers only were added).

**Fig 2:**
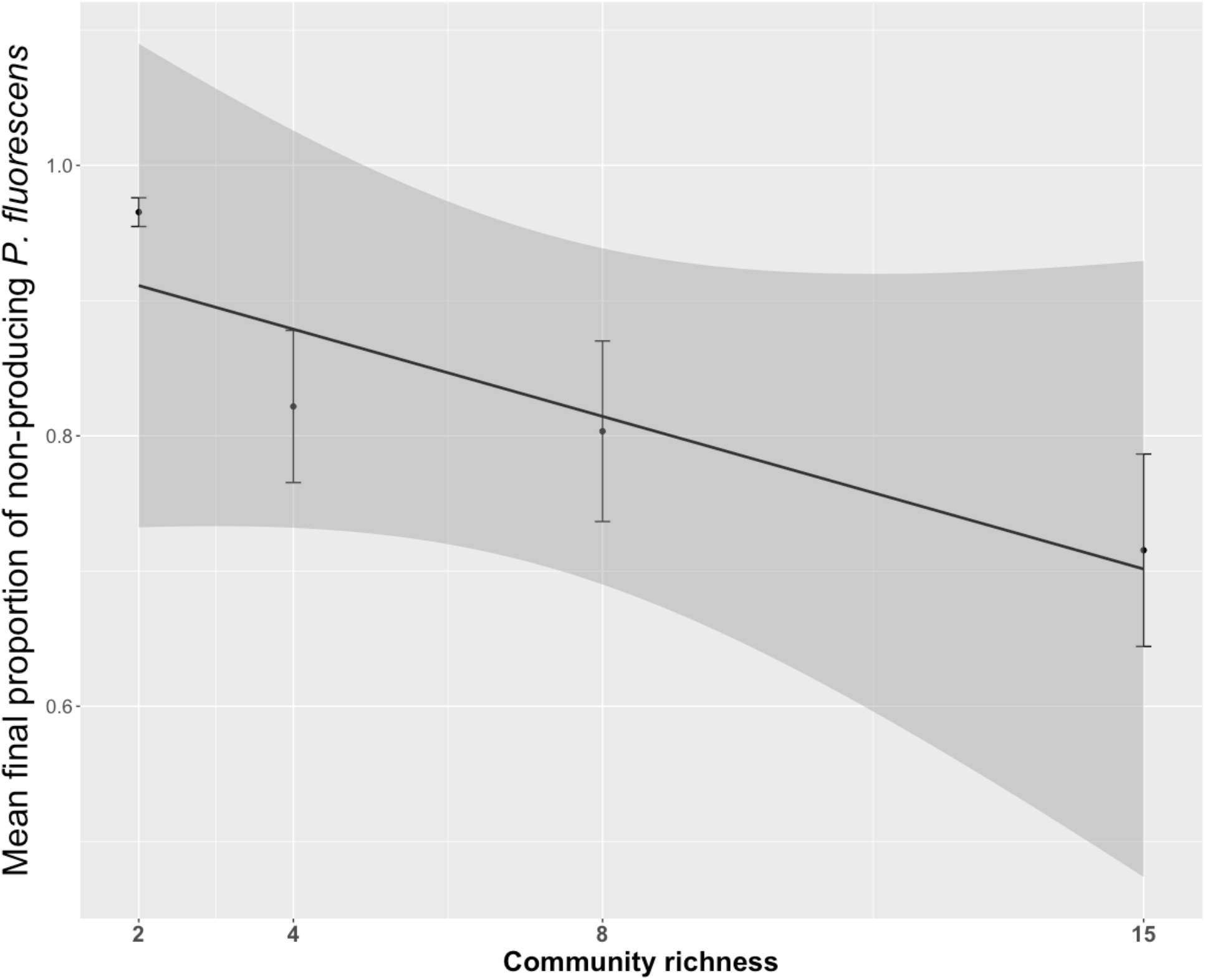
Mean proportion of non-producers in mixed genotype treatments at increasing richness levels. We find that the proportion of non-producers in the community reduced as species richness increased (*X*^2^_1,2_= 10.27, p=0.001). Sample size varies between richness levels. When r=2, n=75. When r=4,8 or 15, n=25. Points show mean value ±SE.

Notably, while the proportion of non-producers decreases with increasing richness, non-producers are never outcompeted by producers – even at species richness 15 – where non-producers still represent 80% of the population. This suggests that although cooperation is relatively more beneficial in a complex rather than simple community, non-producers are still able to exploit producers in all cases.

### c) Disentangling the effect of species richness versus composition on *P. fluorescens* producer and non-producer final cell densities

The above analyses revealed opposing effects of species richness on *P. fluorescens* producer and non-producer populations. However, the above models could not differentiate between the effect of richness *per se* or the added chance of including a biologically important species at high richness levels. Our replication of different compositions at each species richness level allowed us to further investigate the relative roles of richness versus composition on final cell densities [26].

We first investigated whether any particular species had stronger effects than average on final cell densities of our focal *P. fluorescens* producer and non-producer strains growing as single genotypes. In producer populations, species richness and species identity explained 10% and 18% of the variance in final population densities, respectively (Table S2). In non-producer populations, species richness and identity explained 4% and 13% of the variance in final population densities, respectively (Table S3). Together, this suggests that community species composition (rather than species richness *per se)* is an important driver of siderophore dynamics in our focal species.

Our random partition design allowed us to next identify key community species that either promoted or constrained the growth of our focal strain. Linear model coefficients provide the estimated contribution of each species to *P. fluorescens* final densities, relative to an average species’ contribution [26]. We identified one species, sp. 5 *(Erwina sp.)*, that was associated with higher *P. fluorescens* producer and non-producer final densities relative to an average community species (producer: t=2.19, df=150, p<0.05, non-producer: t = 2.21, df=150, p<0.05; Fig 3, Tables S2, S3). However, the positive effect of *Erwina sp*. was weaker for non-producers compared to producers (*Erwina sp*. linear model coefficient for non-producers versus producers was 5.6 and 9.7, respectively), suggesting *Erwina sp*. could tip the balance of intraspecific competition in favour of producers. Furthermore, *Rhodococcus sp (sp8)* was associated with lower densities of non-producers (t = −2.22, df=150, p<0.05), yet had no detectible effect on producers. Together, these findings suggest that the increased likelihood of encountering both *Erwina sp*. and *Rhodococcus sp* at high richness levels can explain the reduction in non-producer densities (relative to producers) growing as single genotypes in diverse versus simple communities.

**Fig 3:**
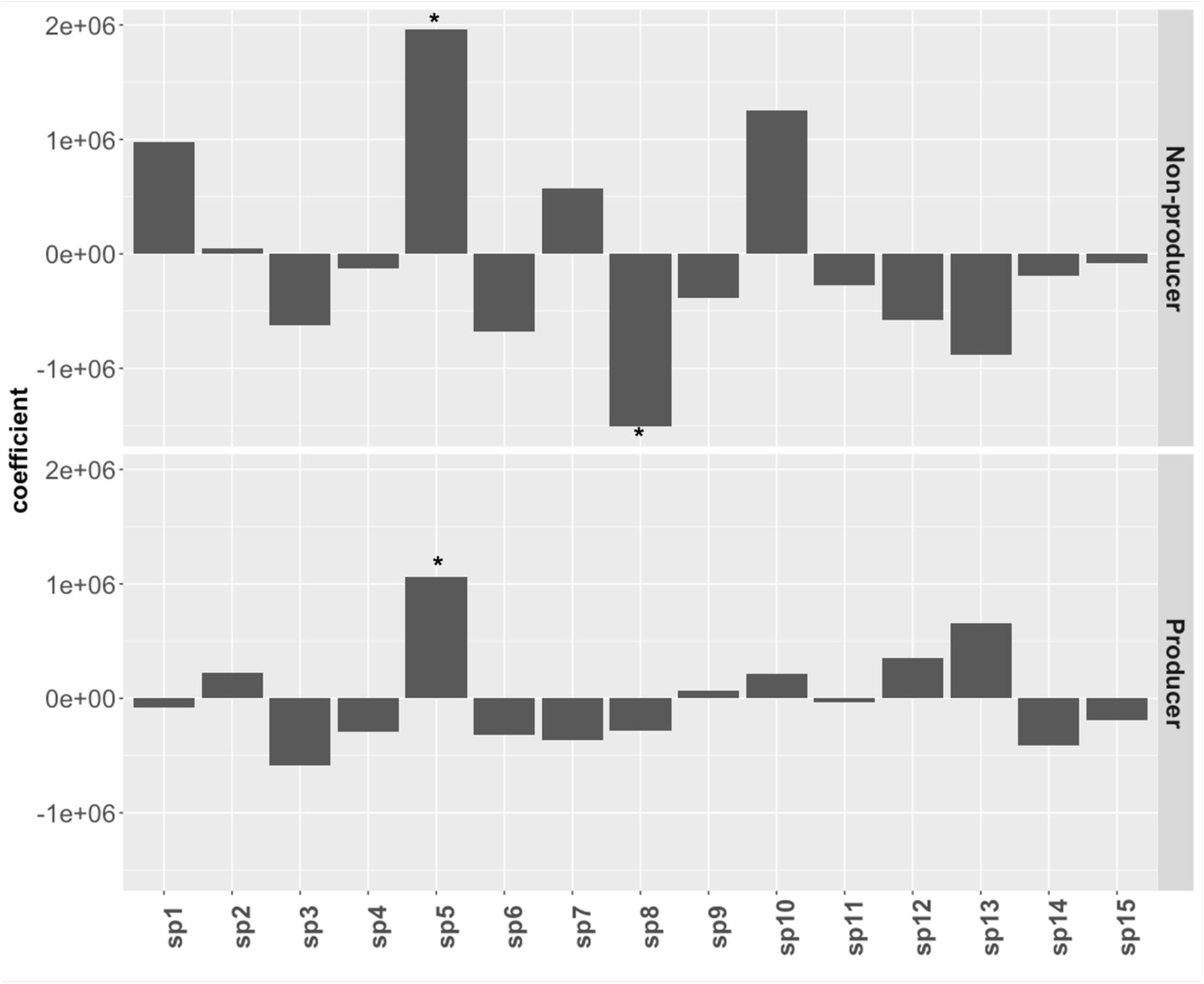
Linear model coefficients for *P. fluorescens* siderophore non-producers (top panel) and producers (bottom panel) strains grown as single genotypes. Positive or negative coefficients indicate species that contribute more or less to *P. fluorescens* growth than average [26]. We find that sp5 (*Erwina sp*) increases final densities of siderophore-producing and non-producing *P. fluorescens*, while sp 8 (*Rhodococcus sp*.) reduces final densities of non-producers only (see main text for statistics).

## 4. Discussion

Here, we empirically test whether the costs and benefits of siderophore production in *P. fluorescens* depend on the composition and richness of the background microbial community. We hypothesised that increased competition for iron in species-rich communities [13, 27] should increase costs of cooperation and consequently favour cheating. However, we find the opposite - cheating is instead constrained in species-rich communities because there is an increased likelihood of encountering species that limit the growth of nonproducers when diversity is high. Our findings suggest that in natural microbial communities, maintaining a diverse microbiota may contribute to the maintenance of siderophore cooperation.

Our finding that siderophore exploitation did not favour non-producers in more diverse communities was surprising, since increased competition for resources in diverse communities are predicted to increase costs of cooperation and hence benefits of cheating [28]. In addition to our finding that cheat-suppressing species could promote cooperation, other ecological conditions could also have contributed to this effect. Firstly, cell counts were quantified after 7 days growth in batch culture. Siderophore producers are less exploitable in stationary phase [29] (since production itself is downregulated in late exponential and stationary growth phases), and species within more diverse communities tend to reach stationary phase earlier [30]. This implies that opportunities for siderophore exploitation (i.e. during mid-exponential phase) are reduced in species rich communities. Secondly, initial cell densities of *P. fluorescens* are higher in low diversity communities due to the nature of our design (i.e starting total cell densities were equal for all communities). Within-species exploitation is greater at high cell densities, since cheats are better able to exploit producers when they are physically closer to them [31]. Hence, our low species richness communities (which were initiated with higher densities of *P. fluorescens* compared to high richness communities), potentially created more opportunities for siderophore exploitation compared to species rich communities. Ultimately, cooperation in communities is likely to be driven by complex interactions between biotic and abiotic factors, where the ecological effects of a community on a species’ cell densities and growth rates are experienced alongside the effects of species interactions themselves.

Our experiments support the idea that genotypic variation within a focal strain (in this case, a single SNP in *pvdL*) can markedly shape the nature of interactions between species in a community [32]. Species interactions in a community can in turn alter the outcome of competition between genotypes within a species – especially when interacting species have genotypic-specific effects [32]. Our study identified a key species in the community (*Rhodococcus sp*) that shaped the growth of our two *P. fluorescens* genotypes in opposing ways. Hence, interspecific interactions in communities can have important consequences for within-species dynamics. We had no prior hypotheses for which species should particularly influence the relative performance of producers and non-producers, and future work would be needed to determine why these two species in particular had these effects. We note, however, that species of *Rhodococcus* [33] produces siderophores, allowing colonization of iron-poor rocky habitats [34]. We hypothesize that *Rhodococcus* might outcompete *P. fluorescens* non-producers for iron using siderophores that are not accessible for uptake by *P. fluorescens* non-producers.

In 2-species communities, we find that non-producers reach higher final densities than producers when grown as single genotypes – where non-producers could not access conspecific siderophores. While this is a surprising result, we pose two possible explanations. Firstly, while the benefits of iron-chelating siderophores tend to be species-specific (due to the requirement of a species specific siderophore receptor), there has been some evidence of siderophore piracy occurring between species. Hence, non-producers may be able to exploit siderophores produced by the second species. Indeed many of our community species are well-known to produce siderophores, such as *Acinetobactor sp*. [35], *Bacillus sp*. [36], *Pantoea sp*. [37], *Serratia sp*. [38], *Pseudomonas sp*. [39] and *Aeromonas sp*. [40]. However, the rarity of inter-species siderophore exploitation in nature coupled with our finding that the cost of cheating is negated in the presence of every single community species makes this unlikely. Secondly, there is some evidence that dead cells can be used as a source of iron for growing populations [15]. Since our final cell counts were taken after 7 days (i.e. when populations were in late stationary phase), cell death could have played a significant role in allowing non-producers to access iron. While further work will be needed to elucidate the mechanism behind this finding its clear that the assumed high cost of cheating in a single species environment may not be observable in a natural community.

Finally, although we are limited here to just 15 species and one community type (tree hole) our results pose interesting questions about the role of the microbiota in maintaining public goods production. Siderophore production is key for promoting plant growth in *P. fluorescens* as a biocontrol agent against plant pathogens [19] yet cheats abound in nature that can compromise its efficacy. It is possible that maintaining a healthy diverse soil microbiota could constrain the fixation of cheats. More generally, the production of public goods such as siderophores, antibiotic resistance via excreted detoxifying systems, and secreted virulence factors have important implications for patient health, ecosystem services and biotechnology. It is therefore important to understand how these systems are affected by the presence of additional non-focal taxa in naturally diverse microbiomes.

