## Supplementary material for "Public goods exploitation is reduced in species-rich microbial communities"

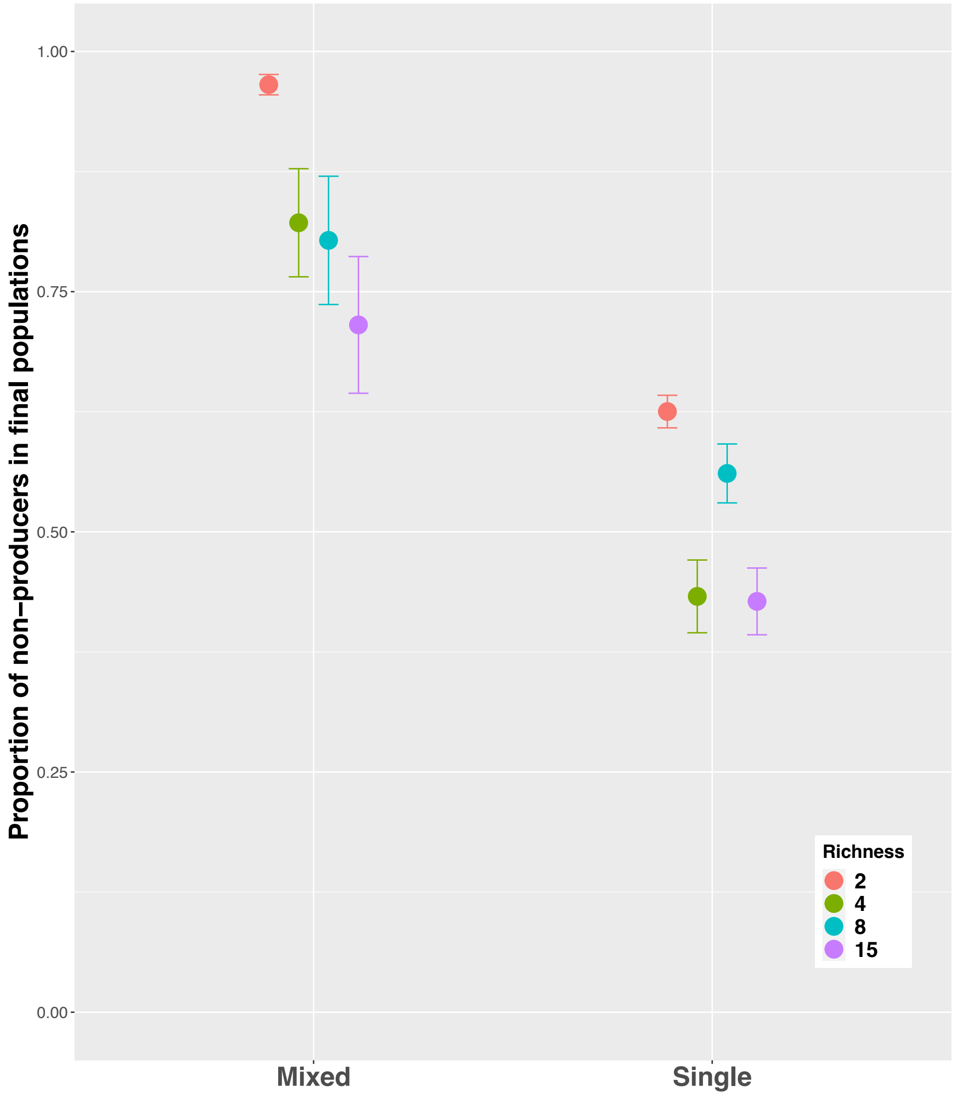


**Supplementary Figure S1.** Final proportion of non-producers to producers of *P. fluorescens* comparing treatments where they grew together in a mixture (mixed genotype treatment) or separately (single genotype treatment). Mean and standard errors for different species richness levels are shown. Non-producers reached higher frequencies when inoculated in the presence of a producer (i.e. mixed genotype treatments). This effect was independent of community richness. See main text for statistics.

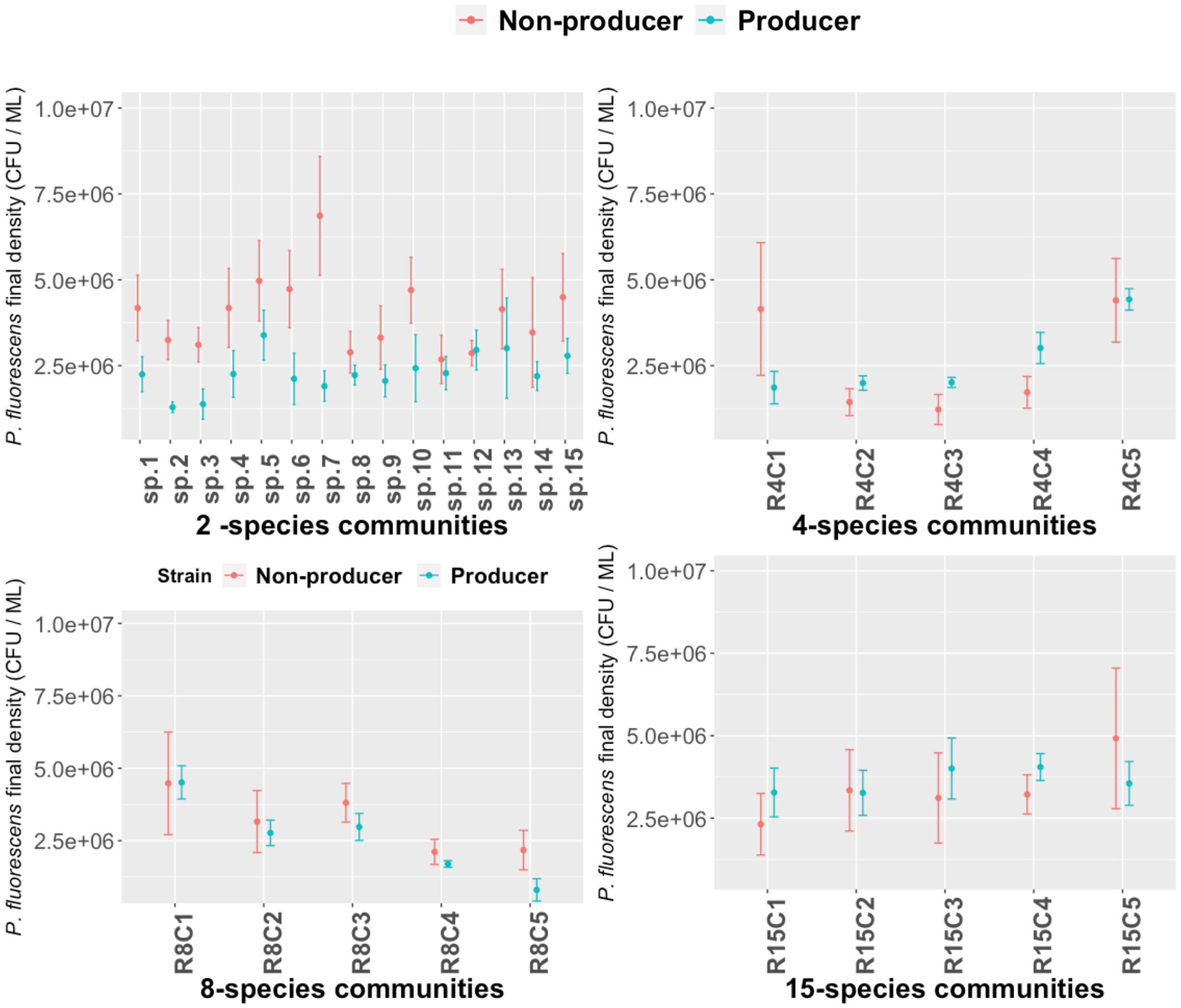


**Supplementary Figure S2.** *P. fluorescens* final densities (CFU/ ML) after growing individually as single genotypes, in communities ranging from 2-15 species. We find significantly greater (173%) non-producer (orange) versus producer (blue) final densities in two-species communities, but no difference between producer and non-producer final cell densities in 4, 8 and 15 species communities (see main text for statistics). Mean and standard errors for different species richness levels are shown.

### Supplementary Methods

**16S sequencing**: DNA was extracted from the twelve isolates using a sodium dodecyl sulphate (SDS) extraction method: A loop of bacteria from culture was suspended in 0.5% SDS; the isopropanol-precipitated DNA was pelleted by spinning at 13,100 rpm for 10 min in a centrifuge, and the pellet was washed in ethanol and re-suspended in 50 µl of sterile water. All chemicals used were obtained from VWR International, except proteinase K and cetrimonium bromide (Sigma-Aldrich). PCR-amplification of 1,500 bp fragments of 16S rDNA was performed using oligonucleotide primers 27F (5'-AGAGTTTGATCMTGGCTCAG- 3') and 1492R (5'-TACGGYTACCTTGTTACGACTT-3') (InvitrogenTM, Thermo Fisher Scientific), using the Mastercycler® gradient thermo-cycler (Eppendorf). The PCR reaction mix of 25µl per sample contained 2µl of template DNA, 12.5µl of RedTaq®, 1.5µl of each primer and 7.5µl of molecular-grade water (Sigma-Aldrich). Reaction mixes were loaded into the thermo-cycler to amplify fragments. PCR products were verified using gel electrophoresis. The 16S rDNA amplicons from 12 isolates were sent to Macrogen© for Sanger sequencing (Sanger et al., 1977). Forward and reverse sequences were aligned using the bioinformatics software Geneious© (Biomatters) and the consensus sequences were manually cleaned in a conservative manner to ensure accuracy. The consensus sequences were then used to identify the isolates via BLAST (Altschul et al., 1990) and using the Sequence Match function in Ribosomal Database Project (Cole et al., 2014).

Altschul, S.F., Gish, W., Miller, W., Myers, E.W., Lipman, D.J. (1990) Basic local alignment search tool. Journal of molecular biology. 215(3), 403–10.

Cole, J.R., Wang, Q., Fish, J. a, Chai, B., McGarrell, D.M., Sun, Y., Brown, C.T., Porras- Alfaro, A., Kuske, C.R., Tiedje, J.M. (2014) Ribosomal Database Project: data and tools for high throughput rRNA analysis. Nucleic acids research. 42(Database issue), D633–42.

**Table S1:** Using a random partitions design (see [26]), we tested whether the growth of our focal *P. fluorescens* producer and non-producer was affected by the richness of the background community. Four levels of community richness: 2, 4 ,8, and 15 was tested. Each richness level includes the addition of *P. fluorescens*, for example, a community richness of 4 represents *P. fluorescens* plus 3 other species. Each richness level was represented by 5 random combinations of the 15 bacterial isolates, except when richness was equal to two; when equal to two, all 15 background isolates were grown with *P. fluorescens*. Each community was replicated five times.

| Richness | Community ID | Species included |
|---|---|---|
| **2** | Sp.1 | 1 |
| **2** | Sp.2 | 2 |
| **2** | Sp.3 | 3 |
| **2** | Sp.4 | 4 |
| **2** | Sp.5 | 5 |
| **2** | Sp.6 | 6 |
| **2** | Sp.7 | 7 |
| **2** | Sp.8 | 8 |
| **2** | Sp.9 | 9 |
| **2** | Sp.10 | 10 |
| **2** | Sp.11 | 11 |
| **2** | Sp.12 | 12 |
| **2** | Sp.13 | 13 |
| **2** | Sp.14 | 14 |
| **2** | Sp.15 | 15 |
| **4** | R4C1 | 1, 4, 11 |
| **4** | R4C2 | 3, 6, 9 |
| **4** | R4C3 | 6, 7, 8 |
| **4** | R4C4 | 4, 8, 9 |
| **4** | R4C5 | 2, 10, 12 |
| **8** | R8C1 | 2, 5, 7, 9, 11, 13, 15 |
| **8** | R8C2 | 1, 3, 5, 8, 9, 12,14 |
| **8** | R8C3 | 2, 3, 4, 7, 10, 11, 13 |
| **8** | R8C4 | 1, 4, 6, 7, 13, 14, 15 |
| **8** | R8C5 | 3, 4, 7, 8, 10, 12, 15 |
| **15** | R15C1 | 2, 3, 4, 5, 6, 7, 8, 9, 10, 11, 12, 13, 14, 15 |
| **15** | R15C2 | 1, 2, 4, 5, 6, 7, 8, 9, 10, 11, 12, 13, 14, 15 |
| **15** | R15C3 | 1, 2, 3, 4, 5, 6, 8, 9, 10, 11, 12, 13, 14, 15 |
| **15** | R15C4 | 1, 2, 3, 4, 5, 6, 7, 8, 10, 11, 12, 13, 14, 15 |
| **15** | R15C5 | 1, 2, 3, 4, 5, 6, 7, 8, 9, 10, 11, 12, 14, 15 |

**Table S2:** ANOVA summary table for linear models partitioning the variance between species ID and community richness in explaining siderophore producer final cell densities. For full details on the random partitions design, see [26].

| | Richness | | | | | Resid(ID) | | | | |
|---|---|---|---|---|---|---|---|---|---|---|
| *Predictors* | *Estimates* | *std. Error* | *CI* | *Statistic* | *p* | *Estimates* | *std. Error* | *CI* | *Statistic* | *p* |
| (Intercept) | 2302466.67 | 165526.86 | 1975328.37 – 2629604.96 | 13.91 | **<0.001** | | | | | |
| richness [4] | 360733.33 | 331053.72 | -293543.25 – 1015009.92 | 1.09 | 0.278 | | | | | |
| richness [8] | 249693.33 | 331053.72 | -404583.25 – 903969.92 | 0.75 | 0.452 | | | | | |
| richness [15] | 1328293.33 | 331053.72 | 674016.75 – 1982569.92 | 4.01 | **<0.001** | | | | | |
| sp1 [0] | | | | | | 108861.29 | 160843.74 | -209259.60 – 426982.17 | 0.68 | 0.500 |
| sp1 [1] | | | | | | -83213.56 | 381213.52 | -837187.49 – 670760.36 | -0.22 | 0.828 |
| sp2 [1] | | | | | | 224439.89 | 443666.30 | -653054.75 – 1101934.54 | 0.51 | 0.614 |
| sp3 [1] | | | | | | -582197.78 | 313471.69 | -1202190.16 – 37794.61 | -1.86 | 0.065 |
| sp4 [1] | | | | | | -295471.22 | 374698.30 | -1036559.16 – 445616.72 | -0.79 | 0.432 |
| sp5 [1] | | | | | | 1059442.29 | 483284.98 | 103588.80 – 2015295.77 | 2.19 | **0.030** |
| sp6 [1] | | | | | | -320442.87 | 333231.51 | -979516.74 – 338631.01 | -0.96 | 0.338 |
| sp7 [1] | | | | | | -367970.57 | 342552.65 | -1045480.02 – 309538.89 | -1.07 | 0.285 |
| sp8 [1] | | | | | | -281327.53 | 369599.06 | -1012330.07 – 449675.01 | -0.76 | 0.448 |
| sp9 [1] | | | | | | 66770.65 | 330332.86 | -586570.19 – 720111.49 | 0.20 | 0.840 |
| sp10 [1] | | | | | | 215823.59 | 457578.49 | -689186.93 – 1120834.11 | 0.47 | 0.638 |
| sp11 [1] | | | | | | -35999.97 | 412718.92 | -852286.08 – 780286.14 | -0.09 | 0.931 |
| sp12 [1] | | | | | | 351350.61 | 445537.34 | -529844.62 – 1232545.83 | 0.79 | 0.432 |
| sp13 [1] | | | | | | 660308.51 | 388494.66 | -108066.24 – 1428683.25 | 1.70 | 0.092 |
| sp14 [1] | | | | | | -411957.96 | 454864.08 | -1311599.86 – 487683.94 | -0.91 | 0.367 |
| sp15 [1] | | | | | | -191790.57 | 400823.42 | -984549.46 – 600968.32 | -0.48 | 0.633 |
| Observations | 150 | | | | | 150 | | | | |
| $R^2$ / $R^2$ adjusted | 0.100 / 0.081 | | | | | 0.179 / 0.081 | | | | |

**Table S3:** ANOVA summary table for linear models partitioning the variance between species ID and community richness in explaining siderophore non-producer final cell densities. For full details on the random partitions design, see [26].

| | Richness | | | | | Resid(ID) | | | | |
|---|---|---|---|---|---|---|---|---|---|---|
| *Predictors* | *Estimates* | *std. Error* | *CI* | *Statistic* | *p* | *Estimates* | *std. Error* | *CI* | *Statistic* | *p* |
| (Intercept) | 3990613.33 | 288727.58 | 3419987.84 – 4561238.83 | 13.82 | **<0.001** | | | | | |
| richness [4] | -1401533.33 | 577455.16 | -2542784.32 – -260282.34 | -2.43 | **0.016** | | | | | |
| richness [8] | -840693.33 | 577455.16 | -1981944.32 – 300557.66 | -1.46 | 0.148 | | | | | |
| richness [15] | -608093.33 | 577455.16 | -1749344.32 – 533157.66 | -1.05 | 0.294 | | | | | |
| sp1 [0] | | | | | | 349075.55 | 295398.45 | -235171.11 – 933322.21 | 1.18 | 0.239 |
| sp1 [1] | | | | | | 981120.26 | 700119.79 | -403594.69 – 2365835.21 | 1.40 | 0.163 |
| sp2 [1] | | | | | | 49546.11 | 814817.79 | -1562021.50 – 1661113.72 | 0.06 | 0.952 |
| sp3 [1] | | | | | | -622941.48 | 575708.16 | -1761591.90 – 515708.94 | -1.08 | 0.281 |
| sp4 [1] | | | | | | -129805.27 | 688154.21 | -1490854.39 – 1231243.86 | -0.19 | 0.851 |
| sp5 [1] | | | | | | 1959960.60 | 887579.69 | 204482.63 – 3715438.57 | 2.21 | **0.029** |
| sp6 [1] | | | | | | -677877.12 | 611998.17 | -1888302.86 – 532548.62 | -1.11 | 0.270 |
| sp7 [1] | | | | | | 570299.46 | 629116.96 | -673984.26 – 1814583.18 | 0.91 | 0.366 |
| sp8 [1] | | | | | | -1509219.68 | 678789.18 | -2851746.40 – -166692.96 | -2.22 | **0.028** |
| sp9 [1] | | | | | | -382743.31 | 606674.63 | -1582640.02 – 817153.39 | -0.63 | 0.529 |
| sp10 [1] | | | | | | 1254071.33 | 840368.29 | -408030.71 – 2916173.38 | 1.49 | 0.138 |
| sp11 [1] | | | | | | -272179.74 | 757981.20 | -1771334.48 – 1226975.00 | -0.36 | 0.720 |
| sp12 [1] | | | | | | -577883.59 | 818254.05 | -2196247.52 – 1040480.35 | -0.71 | 0.481 |
| sp13 [1] | | | | | | -885837.12 | 713492.00 | -2296999.98 – 525325.73 | -1.24 | 0.217 |
| sp14 [1] | | | | | | -188109.03 | 835383.13 | -1840351.31 – 1464133.24 | -0.23 | 0.822 |
| sp15 [1] | | | | | | -81067.72 | 736134.46 | -1537013.41 – 1374877.98 | -0.11 | 0.912 |
| Observations | 150 | | | | | 150 | | | | |
| $R^2$ / $R^2$ adjusted | 0.044 / 0.025 | | | | | 0.130 / 0.026 | | | | |
